# Modulating effects of *FGF12* variants on Na_V_1.2 and Na_V_1.6 associated with Developmental and Epileptic Encephalopathy and Autism Spectrum Disorder

**DOI:** 10.1101/2021.12.03.471090

**Authors:** Simone Seiffert, Manuela Pendziwiat, Tatjana Bierhals, Himanshu Goel, Niklas Schwarz, Amelie van der Ven, Christian Malte Boßelmann, Marjolein H. Willemsen, Ulrike B.S. Hedrich, Ingo Helbig, Yvonne G. Weber

## Abstract

**Objective:** Fibroblast growth factor 12 (FGF12) may represent an important modulator of neuronal network activity and has been associated with developmental and epileptic encephalopathy (DEE). We sought to identify the underlying pathomechanism of FGF12-related disorders.

**Methods:** Patients with pathogenic variants in *FGF12* were identified through published case reports, GeneMatcher and whole exome sequencing of own case collections. The functional consequences of two missense variants and two copy number variants (CNVs) were studied by co-expression of wild-type and mutant FGF12 in neuronal-like cells (ND7/23) with the sodium channels Na_V_1.2 or Na_V_1.6, including their functional active beta-1 and beta-2 sodium channel subunits (*SCN1B* and *SCN2B*).

**Results:** Four variants in *FGF12* were identified for functional analysis: one novel *FGF12* variant in a patient with autism spectrum disorder and three variants from previously published patients affected by developmental and epileptic encephalopathy (DEE). We demonstrate the differential regulating effects of wildtype and mutant FGF12 on Na_V_1.2 and Na_V_1.6 channels. Here, *FGF12* variants lead to a complex kinetic influence on Nav1.2 and Nav 1.6, including loss- as well as gain-of function changes in fast inactivation as well as loss-of function changes in slow inactivation.

**Interpretation:** For the first time, we could demonstrate the detailed regulating effect of FGF12 on Na_V_1.2 and Na_V_1.6 and confirmed the complex effect of FGF12 on neuronal network activity. Our findings expand the phenotypic spectrum related to *FGF12* variants and elucidate the underlying pathomechanism. Specific variants in FGF12-associated disorders may be amenable to precision treatment with sodium channel blockers.

## Introduction

Voltage-gated sodium channels (Na_V_s) play an important role in the initiation and propagation of action potentials in neurons. Na_V_s are expressed in the central and the peripheral nervous system, the skeletal muscle and the heart.^1^ A total of nine different alpha subunits are known, of which variants in five of these subunits (Na_V_1.1 [*SCN1A*], Na_V_1.2 [*SCN2A*], Na_V_1.3 [*SCN3A*], Na_V_1.6 [*SCN8A*] and Na_V_1.7 [*SCN9A*]) have been related to epilepsy or intellectual disability.^2–4^ Sodium channels are modulated by different proteins. Of these, four members of the FGF family also called FGF11 subfamily, namely FGF11, FGF12, FGF13 and FGF14, play a major role in modulating sodium channels activation and fast inactivation kinetics.^5^ All members of the FGF11 subfamily have two transcripts that differ in their N-terminal length, suggesting that they are involved in different molecular mechanisms (Fig 1A).^6^ Accordingly, they exert a broad spectrum of modulating effects on various channels. FGF14 differentially modulates the fast inactivation of Na_V_1.2 and Na_V_1.6, and slows the recovery from inactivation of Na_V_1.6.^7^ FGF13 is able to modulate Na_V_1.5 by influencing fast inactivation as well as the recovery of fast inactivation.^8^ FGF12 is known to interact with Na_V_1.2, Na_V_1.5 and Na_V_1.6, while it seems not to modulate Na_V_1.1.^9–12^

**Figure 1:**
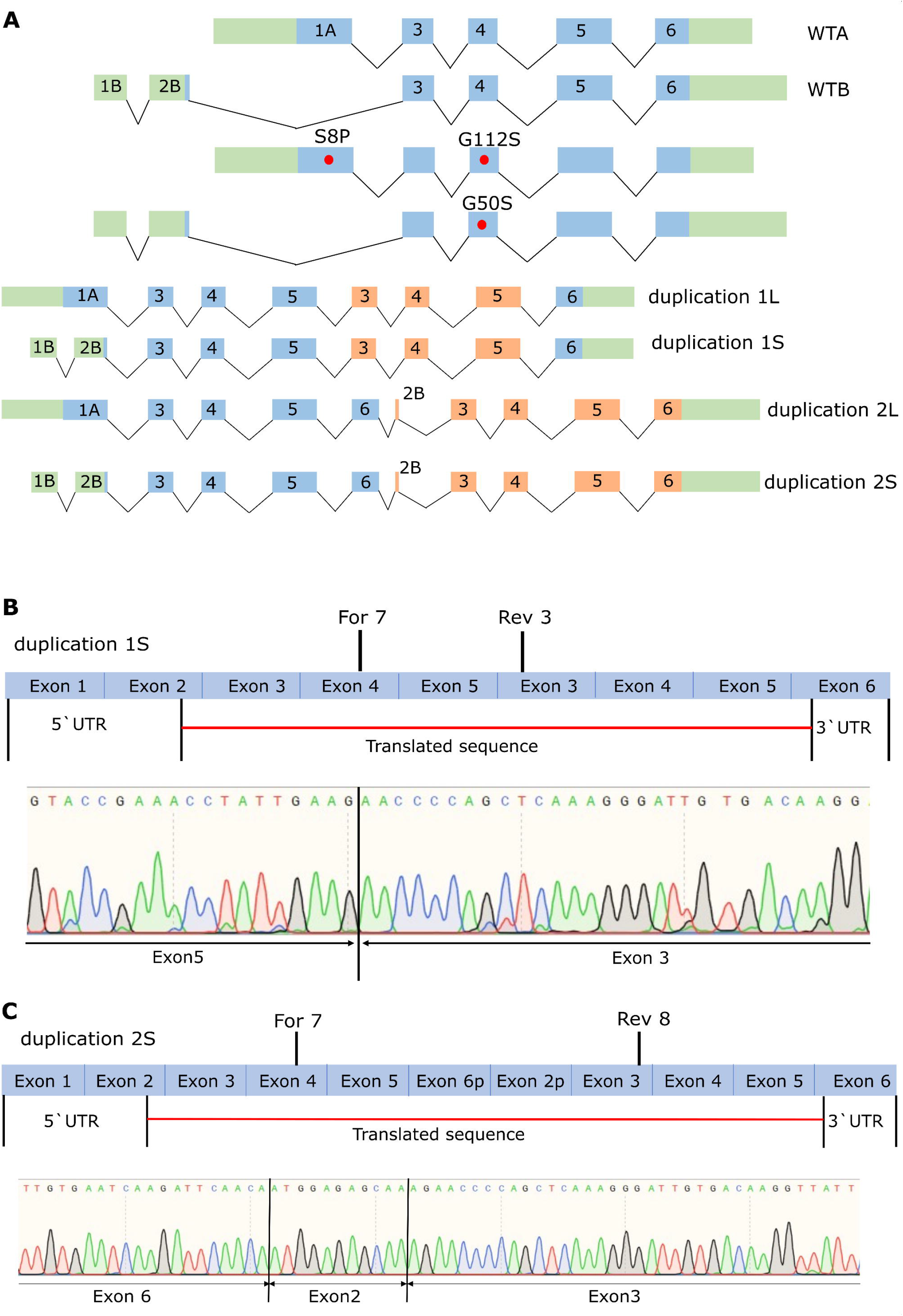
Characterisation of *FGF12* CNVs. A. Schematic drawings of the different FGF12 WTs as well as the different missense variants and CNVs. Green: UTR sites, blue: translated exons, orange: duplicated translated exons, red dots: missense variants B. Characterisation of duplication 1 using RT-PCR from cDNA extracted from venous blood of the two patients described in Willemsen et al., 2020. Primer For 7 and Rev 3 were used for the amplification of the CNV, and the sequencing result of this PCR product showed a duplication encompassing Exon 3 to Exon 5. C. Characterisation of duplication 2 using RT-PCR from RNA extracted from Fibroblasts of the patient described in Willemsen et al., 2020. Primer For 7 and Rev 8 were used for the amplification of the CNV, and the sequencing result of this PCR product showed a duplication encompassing the whole gene of FGF12 (Exon 2 to Exon 6).

Variants in genes encoding members of this subfamily have been associated with generalized epilepsy with febrile seizures plus (GEFS+; FGF13), spinocerebellar ataxia (FGF14) or developmental and epileptic encephalopathy (DEE; FGF12).^13–15^ However, the detailed pathophysiological mechanism behind FGF12-associated DEE has been unclear.^16^

Here we investigated the modifying effects of FGF12 on Nav1.2 and Nav1.6 in the wildtype state. Additionally, we carried out functional analyses of four *FGF12* variants, including a novel variant of *FGF12* causing autism spectrum disorder (ASD).

## Patients and Methods

### Patients and variants collection

The variants were collected through GeneMatcher reports^17^, literature review and whole exome sequencing in two novel cases identified by the Institute of Human genetics UKE Hamburg-Eppendorf and the Clinic for Neuropediatric UKSH Kiel. Further pathogenic variants were not detected based on the classification criteria of the American College of Medical Genetics^18^. Segregation analysis was performed if applicable. For the novel cases, written informed consent was obtained from all participants or their legal representatives. The current study was approved by each local ethics committee.

### Fibroblast generation

The fibroblasts were generated from a skin biopsy with a size of 3-4 mm. First, the subcutaneous adipose tissue was removed from the biopsy and washed twice with PBS + Penicillin/Streptomycin. Afterwards, the biopsy was cut in 1-millimetre pieces and transferred to a 15 mL tube with 2 mL collagenase solution (collagenase diluted in 2 mL DMEM) and incubated for 3 hours at 37°C. After the incubation, 6 mL DMEM + 20% FCS was added to dilute the collagenase. Once spun down, the medium was exchanged with 6 mL fresh DMEM-medium, supplemented with 20 % FCS. The sample was split into 4 T25 flasks and incubated for 7 days at 37°C in 5% CO_2_. The medium was refreshed every 3-4 days, as cells started to grow.

### Transcript identification

For three duplication cases, we had to determine the structure of the resulting transcript. The duplication 3q28q29(chr3:191.860.089-192.451.114), which was found in one individual, and the duplication 3q28q29 (chr3:191876978-192454675), which was identified in two related individuals, have been published previously within a clinical report.^19^ To analyse the transcript of the first duplication, we performed RNA purification from fibroblasts and retrotranscription to cDNA, using the Isolation II RNA Mini kit (Bioline) and the First strand cDNA synthesis kit for RT-PCR (Roche), respectively, according to manufacturer protocols. For the CNV-specific PCR, the following primers were used: FGF12For7: CTC TCT TCA ATC TAA TTC CCG TGG and FGF12Rev8: GTA GTC GCT GTT TTC GTC CTT GGT C. For the analysis of the second duplication, RNA from venous blood of both patients was isolated and retro-transcribed in cDNA followed by a CNV-specific PCR using the following primers: FGF12For7: CTC TCT TCA ATC TAA TTC CCG TGG and FGF12Rev3: CCT TGT CAC AAT CCC TTT GAG CTG G. All three PCR products were sent for sequencing to LGC Genomics GmbH (Berlin, Germany).

## Functional investigations

### Mutagenesis

The human *FGF12* wildtype (WT) and the variants were purchased from Genescript as g- blocks and cloned with the HiFi DNA Assembly Kit (purchased from NEB) into the pCMV6 Vector (purchased from Origene) together with the two human β_1_- and β_2_-subunits of voltage- gated Na^+^ channels as well as the antibiotic puromycin as selection marker. The four proteins were separated by the P2A, E2A and T2A sequences, respectively, to separate the proteins during translation. Except for the stop codon, the sequence of FG12A corresponded to the Ensembl transcript ENST00000454309.6 and Consensus Coding Sequence Database transcript CCDS3301 and FGF12B corresponded to the Ensembl transcript ENST00000445105.6 and Consensus Coding Sequence Database transcript CCDS46983.

The human Na_V_1.6 channel construct was purchased from Origene and modified to introduce TTX resistance by a known point mutation (c.1112A > G, p.Y371C).^20^ The WT open reading frame included the canonical *SCN8A* coding sequence, with a C-terminal P2A sequence combined with an eGFP. The *SCN8A* coding sequence (5940 bp) contained the splice isoform 5 N of exon 5. Apart from the TTX resistance change and the stop codon, it was identical to the coding sequences of Ensembl transcript ENST00000354534.11 and Consensus Coding Sequence Database transcript CCDS44891. To engineer the TTX-resistance into the human Na_V_1.6 channel, site-directed mutagenesis was performed using PCR with Pfu polymerase (Promega; mutagenic primers are available upon request). To record currents from Na_V_1.2 channels, we exchanged the Na_V_1.6 cDNA with the Na_V_1.2 cDNA and introduced TTX resistance by a known point mutation (c.1154 T>C, p. F385S).^21^ Except for the stop codon and TTX resistance change, the cDNA was identical to the coding sequence of Ensembl transcript ENST00000636071.2 and Consensus Coding Sequence Database transcript CCDS33313. To engineer the TTX-resistance into the human Na_V_1.2 channel, site-directed mutagenesis was performed using PCR with Phusion HF polymerase (NEB; mutagenic primers are available upon request). Further mutations in the constructs were excluded by sequencing the whole open reading frame prior using the clones for physiological experiments.

### Transfection of ND7/23 cells

ND7/23 is a hybrid cell line derived from neonatal rat dorsal root ganglia neurons fused with mouse neuroblastoma cells.^22^ It was purchased from Sigma Aldrich and cultured in Dulbecco’s modified Eagle nutrient medium (Invitrogen) supplemented with 10% fetal calf serum (PAN-Biothech) and 1% L-glutamine 200 mM (Biochrom) at 37°C, with 5%

CO_2_ humidified atmosphere. ND7/23 cells were plated in 35 mm petri dishes following the standard protocol for Lipofectamine™ 3000 (Invitrogen) transfections.

Transfections of WT or mutant human *FGF12* cDNAs were then performed together with the human *SCN8A* cDNA, encoding the Na_V_1.6 channel α-subunit with an engineered TTX resistance or the human *SCN2A* cDNA, encoding the Na_V_1.2 channel α-subunit with an engineered TTX resistance. For co-expression of FGF12 with both β-subunits and the α subunit, 3.4 µg of cDNA were used (3 µg of the α-subunit and 0.4 µg of the *FGF12* with both β-subunits). The amount of the different subunits based on our previous publication on this subject.^3^ After 24 h, cells were selected by adding 1.5 µg/mL puromycin (InvivoGen) to the medium. Electrophysiological recordings were performed 72 h after transfection only from cells expressing all four proteins, which were recognized by (i) a green fluorescence (α- subunit) and (ii) a puromycin resistance (FGF12 and both beta subunits).

### Electrophysiology

For recordings of transfected ND7/23 cells, 500 nM TTX was added to the bath solution to block all endogenous Na^+^ currents. Standard whole-cell voltage clamp recordings were performed using an Axopatch 200B amplifier, a Digidata 1440 A digitizer and Clampex 10.2 data acquisition software (Molecular Devices) as described previously.^3^ Leakage and capacitive currents were automatically subtracted using a prepulse protocol (−P/4). Cells were held at −100 mV. Currents were filtered at 10 kHz and digitized at 20 kHz. Cells were visualized using an inverted microscope (DM IL LED, Leica). All recordings of transfected ND7/23 cells were performed 10 min after establishing the whole-cell configuration at room temperature to avoid larger shifts in voltage dependence. Borosilicate glass pipettes had a final tip resistance of 1.5–3.5 MΩ when filled with internal recording solution (see below). to 90% compensation was always <5 mV. The pipette solution contained (in mM): 10 NaCl, 1 EGTA, 10 HEPES, 140 CsF (pH was adjusted to 7.3 with CsOH, osmolarity was adjusted to 310 mOsm/kg with mannitol). The bath solution contained (in mM): 140 NaCl, 3 KCl, 1 MgCl_2_, 1 CaCl_2_, 10 HEPES, 20 TEACl (tetraethylammonium chloride), 5 CsCl and 0.1 CdCl_2_ (pH was adjusted to 7.3 with CsOH, osmolarity was adjusted to 320 mOsm/kg with mannitol).

### Data recording and statistical analysis

For the voltage-clamp recordings in ND7/23 cells, the biophysical parameters of human WT Na_V_1.6 and human Na_V_1.2 channels were obtained as described previously^3^ with a change of the test pulse of all protocols to 0 mV for the recordings of Na_V_1.2. Furthermore, slow inactivation was determined using 30 s conditioning pulses to various potentials followed by a test pulse to - 10 mV or 0 mV, respectively, at which the peak current reflected the percentage of non-inactivated channels. The activation and fast inactivation protocols were analysed as previously described.^3^ Additionally, for the fast inactivation time constant a single exponential fit was used. For the recovery from fast inactivation the current of each was normalized to its prepulse and then a second-order exponential function was used with an initial delay. The peak amplitude of the first exponential function (A1) was set to 0.3 for all curves. All data were analysed using Clampfit software of pClamp 10.6 (Axon Instruments), Microsoft Excel (Microsoft Corporation, Redmond, WA, USA), or Gnuplot 5.2 (Freeware, T.Williams & C.Kelly). Statistics were performed using Graphpad software (Graphpad prism, San Diego, CA, USA). All data were tested for normal distribution. For comparison of multiple groups, one-way ANOVA with Holm-Sidak’s *post hoc* test was used for normally distributed data and ANOVA on ranks with Dunn’s *post hoc* test was used for not normally distributed data.

## Results

### Patients

We identified two novel patients through our own patient collection and GeneMatcher, both harbouring a *de novo* variant in *FGF12* and suffering from DEE or ASD, respectively.

At the time of recruitment, patient 1 was a 11-year-old boy. His generalized seizures started at three months of age, after which he showed moderate developmental delay without regression, as well asfeatures of ASD. On brain magnetic resonance imaging (MRI), mild global atrophy was noted. Whole exome sequencing revealed a *de novo* variant in *FGF12* (NM_021032.4: c.334G>A, p.G112S), with a known association to DEE (ClinVar accession number: VCV000522854.4).

Patient 2 was a 4.5-year-old boy. At 1.5 years of age, he was diagnosed with ASD. He showed developmental delay with language impairment, feeding difficulties and intellectual disability. ECG, EEG and MRI were normal. Whole exome sequencing revealed a *de novo* variant in the *FGF12* gene (NM_021032.4: c.23C>T, p.S8P), which has not been previously described.

### Variants

In addition to the above-mentioned patients and variants, we included three different copy number variants (CNVs) (3q28q29(191,876,978-192,454,675) X3; 3q28q29(191,876,968- 192,454,685) X3; 3q28q29(191,860,089-192,451,114) X3) which have been published before.^19, 23^ The first duplication was already characterized on a molecular level^23^ while the other two duplications were not further characterized. None of them had been functionally analysed. All of these missense variants and copy number variants were associated with DEE.

### Characterisation of the duplications

To functionally investigate the duplications, it was first necessary to determine the correct resulting transcript form. The first duplication (3q28q29(191.876.978-192.454.675) X3 had already been analysed and published before.^23^ This duplication encompasses Exons 3 to 5 (NM_004113) and is referred to as duplication 1 for the following functional characterization (Fig 1B). The second duplication (3q28q29(191,876,968-192,454,685) X3) is very similar to the first duplication but spans 10 bp more in each direction. To evaluate if this expansion leads to a different transcript, we used reverse transcription PCR from isolated RNA from venous blood of the patient. The 332 bp encompassing PCR product demonstrated Exon 4, 5 followed by Exon 3 (NM_004113) (Fig 1B). Thus, this duplication encompasses the same Exons as duplication 1. The third duplication (3q28q29(191,860,089-192,451,114) X3) was analysed using reverse transcription PCR from isolated RNA of the patient’s fibroblasts. The 542 bp encompassing PCR product showed Exon 4, 5 and 6, which was followed by a part of exon 2 and exon 3 (NM_004113) (Fig 1C). Thus, this duplication (referred here as duplication 2) encompasses the whole gene except for exon 1, partially exon 2 and partially exon 6, since these parts include just the UTR sites.

### Functional characterization of the effect of different FGF12 WT isoforms on Na_V_1.6 and Na_V_1.2 channels

FGF12 has two different homologs which result from alternative splicing and are only different in their N-terminal part (FGF12A, long N-terminal part: WTA; FGF12B, short N-terminal part: WTB) (Fig 1A). Since both homologs include the core structure which is necessary for the interaction of FGF12 with Na_V_s, we characterized the effects of both WTs on Na_V_1.2 and Na_V_1.6. The effect on Na_V_1.5 was not investigated since Na_V_1.5 is not expressed in the brain. Expression of FGF12A together with Na_V_1.6 showed a reduction in Na^+^ current density (Fig 2A and B, Table 1), and altered voltage-dependent gating kinetics including a depolarizing shift of fast inactivation (Fig 2C, Table 1) and slow inactivation (Fig 2D, Table 1). Additionally, fast inactivation and recovery of fast inactivation were slowed (Fig 2E and F, Table 1). Taken together, FGF12A results in a mixed gain- and loss-of overall ion channel function (GOF/LOF) in Na_V_1.6. Conversely, FGF12B leads to a depolarizing shift of fast inactivation as well as to a depolarizing shift of slow inactivation, resulting in a gain-of-function (GOF) effect on Na_V_1.6 (Fig 2C and D, Table 1).

**Figure 2:**
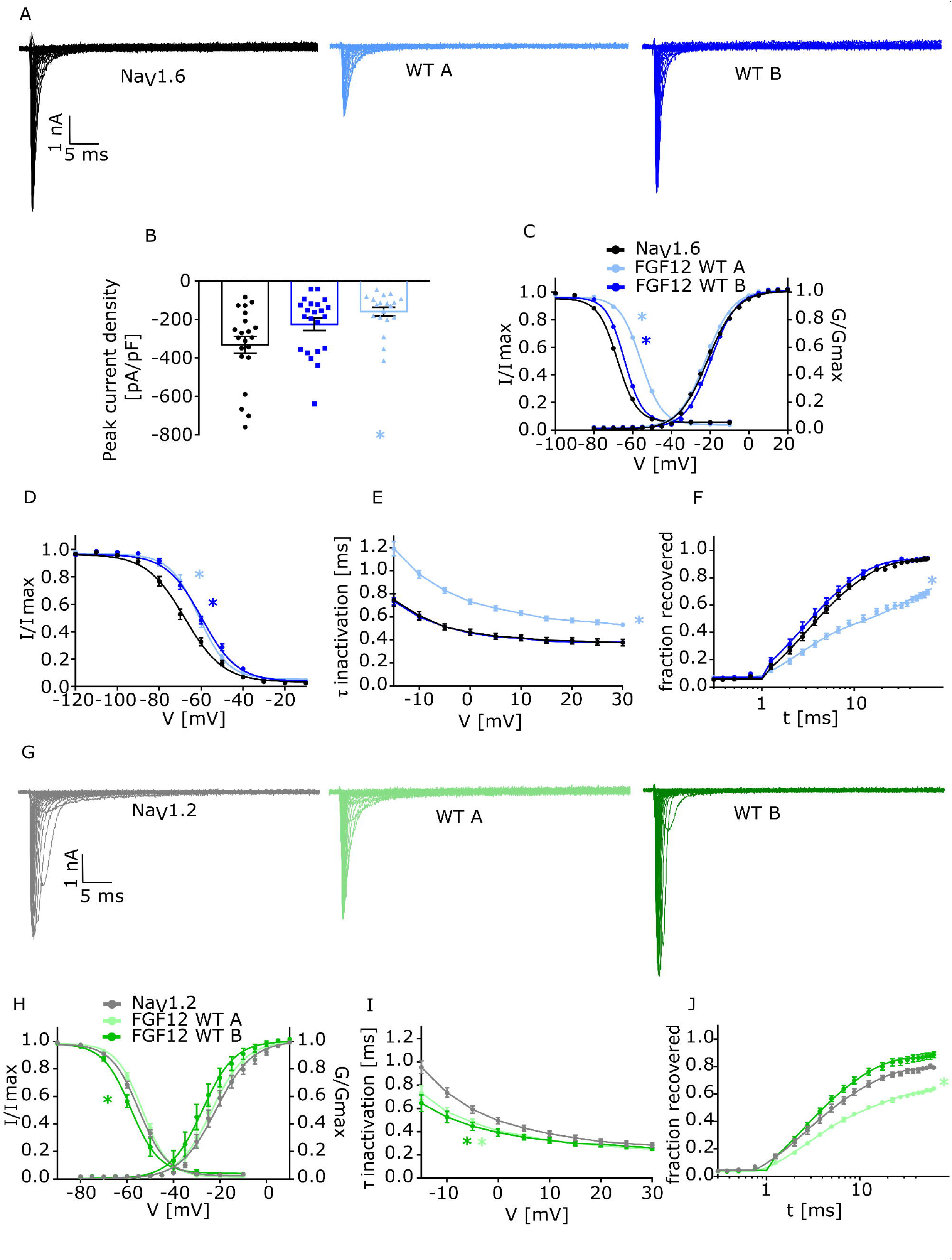
Electrophysiological analysis of the two different FGF12 WT isoforms (WTA and WTB) Functional analysis of the effect of the two different FGF12 WTs on Na_V_1.2 and Na_V_1.6 compared to the channels without FGF12. In summary, the figures show that the expression of WTA together with Na_V_1.6 leads to a mixture of gain- and loss-of function changes while the expression of WTB together with Na_V_1.6 leads to gain-of function changes. Whereas the expression of the two WTs together with Na_V_1.2 leads to loss-of function changes. A. Representative traces of Na_V_1.6 currents in ND7/23 cells expressing Na_V_1.6 without FGF12, with FGF12 WTA or FGF12 WTB in response to voltage steps from -80 to +35 mV in 5 mV steps. B. Mean Current amplitudes of analysed ND7/23 cells injected with Na_V_1.6 without FGF12 (n=21), with FGF12 WTA (n=19) or FGF12 WTB (n=22). Expression of Na_V_1.6 together with FGF12 WTA showed a significantly reduced current density compared to Na_V_1.6 alone. C. Mean voltage-dependent activation and fast inactivation of Na_V_1.6 without FGF12 (n=21), with FGF12 WTA (n=19/16) or FGF12 WTB (n=22). Lines illustrate Boltzmann Function fit to the data points. All fast inactivation curves showed a significant shift to more depolarized potentials in comparison to Na_V_1.6 without FGF12. All data are shown as means ± SEM. D. Mean voltage-dependent slow inactivation of Na_V_1.6 without FGF12 (n=15), with FGF12 WTA (n=11) or FGF12 WTB (n=15). Lines illustrate Boltzmann Function fit to the data points. All slow inactivation curves showed a significant shift to more depolarized potentials in comparison to Na_V_1.6 without FGF12. All data are shown as means ± SEM. E. Mean voltage-dependent fast inactivation time constant of Na_V_1.6 without FGF12 (n=21), with FGF12 WTA (n=19) or FGF12 WTB (n=22). All data are shown as means ± SEM. F. Time course of recovery from fast inactivation of Na_V_1.6 without FGF12 (n=10), with FGF12 WTA (n=10) or FGF12 WTB (n=11) determined at -100 mV. Lines represent fits of biexponential functions yielding the time constants τ1 and τ2. A1 was set to 0.3. All values of electrophysiological results, numbers and p-values are listed in table 1 and are shown as means ± SEM. G. Representative traces of Na_V_1.2 currents in ND7/23 cells expressing Na_V_1.2 without FGF12, with FGF12 WTA or FGF12 WTB in response to voltage steps from -80 to +35 mV in 5 mV steps. H. Mean voltage-dependent activation and fast inactivation of Na_V_1.2 without FGF12 (n=14/13), with FGF12 WTA (n=13) or FGF12 WTB (n=11). Lines illustrate Boltzmann Function fit to the data points. All data are shown as means ± SEM. I. Mean voltage-dependent fast inactivation time constant of Na_V_1.2 without FGF12 (n=14), with FGF12 WTA (n=13) or FGF12 WTB (n=11). All deactivation curves showed a significantly faster deactivation in comparison to channels only containing Na_V_1.2 subunit. All data are shown as means ± SEM. J. Time course of recovery from fast inactivation of Na_V_1.2 without FGF12 (n=12), with FGF12 WTA (n=11) or FGF12 WTB (n=10) determined at -100 mV. Lines represent fits of biexponential functions yielding the time constants τ1 and τ2. A1 was set to 0.3. All values of electrophysiological results, numbers and p-values are listed in table 2 and are shown as means ± SEM.

**Table 1:**
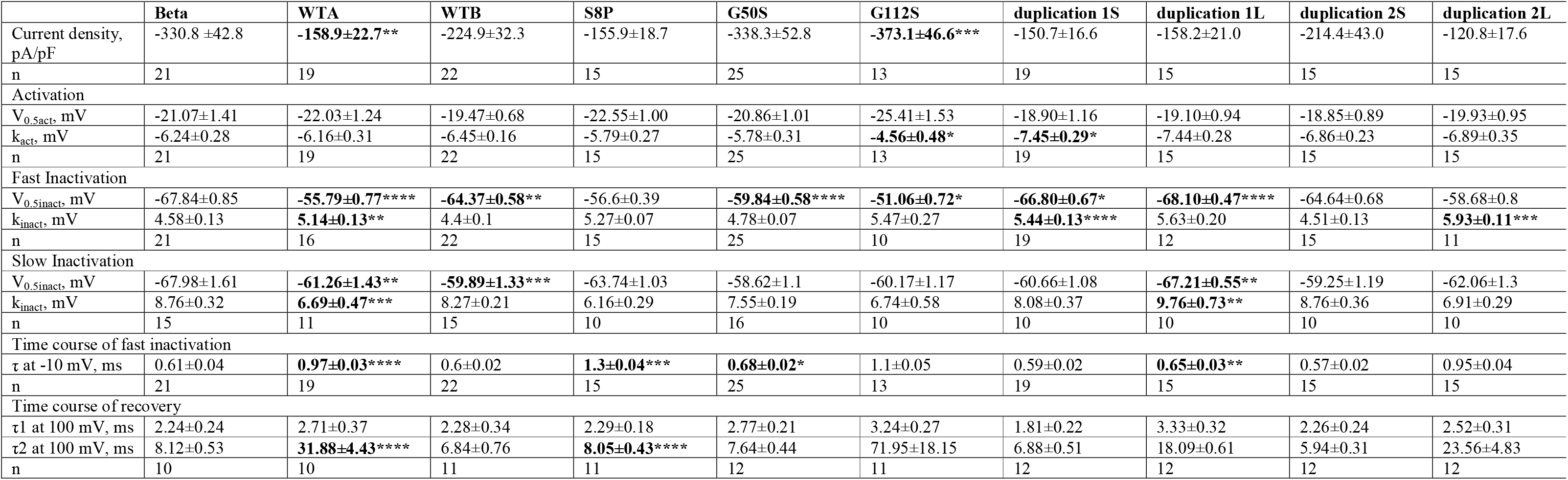
Biophysical parameters of Na_V_1.6 with FGF12 variants. Data are presented as mean ± SEM, k slope factor. Significant differences are shown using *p<0.05, **p<0.005, ***p<,0.0005 ****p<0.0001 and are marked in bold.

**Table 2:** Biophysical parameters of Na_V_1.6 with FGF12 variants. Data are presented as mean ± SEM, k slope factor. Significant differences are shown using *p<0.05, **p<0.005, ***p<,0.0005 ****p<0.0001 and are marked in bold.

|  | Beta | WTA | WTB | S8P | G50S | G112S | duplication 1S | duplication 1L | duplication 2S | duplication 2L |
| --- | --- | --- | --- | --- | --- | --- | --- | --- | --- | --- |
| Current density, pA/pF | -327.58±54.05 | -394.06±58.55 | -349.69±57.98 | -239.94±42.09 | -416.16±76.71 | -257.25±50.24 | -516.67±88.93 | -310.08±86.74 | -496.31±133.62 | -278.73±71.97 |
| n | 13 | 13 | 11 | 11 | 12 | 11 | 12 | 11 | 12 | 12 |
| Activation |  |  |  |  |  |  |  |  |  |  |
| V<sub>0.5act</sub>, mV | -21.25±1.73 | -23.02±1.76 | -26.46±2.39 | -23.00±1.39 | -24.90±1.43 | -21.42±1.58 | -21.32±1.72 | -21.08±3.10 | -23.66±2.53 | -20.52±2.01 |
| k<sub>act</sub>, mV | -6.75±0.51 | -6.18±0.5 | -5.08±0.72 | -6.24±0.57 | -6.05±0.52 | -5.93±0.47 | -6.16±0.41 | -6.24±0.84 | -5.33±0.77 | -6.70±0.50 |
| n | 13 | 13 | 11 | 11 | 12 | 11 | 12 | 11 | 12 | 12 |
| Fast Inactivation |  |  |  |  |  |  |  |  |  |  |
| V<sub>0.5inact</sub>, mV | -54.86±0.72 | -53.35±0.74 | **-57.57±2.03\*** | -51.70±0.63 | -57.62±1.13 | **-48.28±1.01\*** | -58.17±0.69 | **-60.38±0.87\*\*** | -57.96±1.22 | -53.34±0.72 |
| k<sub>inact</sub>, mV | 5.77±0.14 | 5.43±0.08 | **5.20±0.12\*\*** | 5.54±0.14 | **5.74±0.11\*** | 5.51±0.16 | **6.79±0.23\*\*\*\*** | **6.60±0.26\*\*\*\*** | 5.24±0.11 | **6.15±0.13\*\*** |
| n | 13 | 13 | 11 | 10 | 13 | 11 | 11 | 11 | 12 | 12 |
| Slow Inactivation |  |  |  |  |  |  |  |  |  |  |
| V<sub>0.5inact</sub>, mV | -61.55±2.04 | -60.84±1.02 | -59.42±2.13 | -60.39±1.33 | -61.72±1.40 | -57.00±1.21 | -57.45±0.94 | -61.23±1.46 | -57.51±1.06 | -56.68±1.08 |
| k<sub>inact</sub>, mV | 6.29±0.58 | 5.40±0.34 | 6.04±0.59 | 5.61±0.41 | 5.81±0.40 | 5.17±0.24 | 6.65±0.38 | **6.55±0.23\*** | 6.55±0.30 | 5.40±0.22 |
| n | 10 | 10 | 10 | 11 | 12 | 11 | 10 | 10 | 10 | 10 |
| Time course of fast inactivation |  |  |  |  |  |  |  |  |  |  |
| τ at -10 mV, ms | 0.74±0.04 | **0.57±0.04\*** | **0.53±0.05\*\*** | 0.72±0.05 | 0.61±0.03 | **0.77±0.06\*\*** | 0.64±0.05 | 0.61±0.03 | 0.53±0.04 | 0.68±0.04 |
| n | 13 | 13 | 11 | 11 | 12 | 11 | 12 | 11 | 12 | 12 |
| Time course of recovery |  |  |  |  |  |  |  |  |  |  |
| τ<sub>1</sub> at 100 mV, ms | 2.33±0.30 | 2.50±0.18 | 1.90±0.19 | 2.22±0.24 | 2.73±0.23 | 2.81±0.24 | 2.39±0.29 | 2.79±0.23 | 2.16±0.31 | 2.60±0.25 |
| τ<sub>2</sub> at 100 mV, ms | 9.34±0.83 | **17.68±2.10\*\*** | 8.06±0.71 | **10.40±1.12\*** | 10.07±0.68 | 22.80±4.05 | 9.22±0.75 | 13.86±1.27 | 8.51±0.75 | 14.02±1.23 |
| n | 12 | 11 | 10 | 10 | 12 | 10 | 11 | 10 | 11 | 10 |

The effect of FGF12A on Na_V_1.2 was markedly different from the effect on Na_V_1.6. Here, we did not find any change in current density or fast inactivation (Fig 2G and H, Table 2). Similar to Na_V_1.6 channels, FGF12A, when co-expressed with Na_V_1.2, showed an accelerated fast inactivation (Fig 2 I, Table 2) and a slowed recovery from fast inactivation (Fig 2J, Table 2). For co-expression of FGF12B and Na_V_1.2, we found a hyperpolarizing shift of fast inactivation (Fig 2H, Table 2) as well as an accelerated fast inactivation (Fig 2I, Table 2), resulting in an overall loss-of-function (LOF).

### Functional characterization of FGF12 variants on Na_V_1.6 and Na_V_1.2 channels

Previously, we described the FGF12 variant functional effects by co-expression of the variants with either Na_V_1.2 or Na_V_1.6 channels, comparing them to the corresponding WT recordings. The missense variant p.(Ser8Pro) (S8P) is located at the N-terminal part of FGF12A. Since the only difference between FGF12 WTs is the different length of the N-terminal part, the variant S8P can only be found in FGF12 WTA (Fig 1A). The co-expression of S8P with Na_V_1.6 showed no effect on current density or fast inactivation (Fig 3A, B and C, Table 1), but slowed fast inactivation and accelerated recovery from fast inactivation (Fig 3D and E, Table 1). Expression of S8P with Na_V_1.2 demonstrated a similar effect as in Na_V_1.6. Here, no changes in current density or fast inactivation were observed, whereas recovery from fast inactivation was accelerated (Fig 4 A, B, C and D, Table 2). Thus, S8P showed a GOF effect on both Na_V_1.6 and Na_V_1.2 channels.

**Figure 3:**
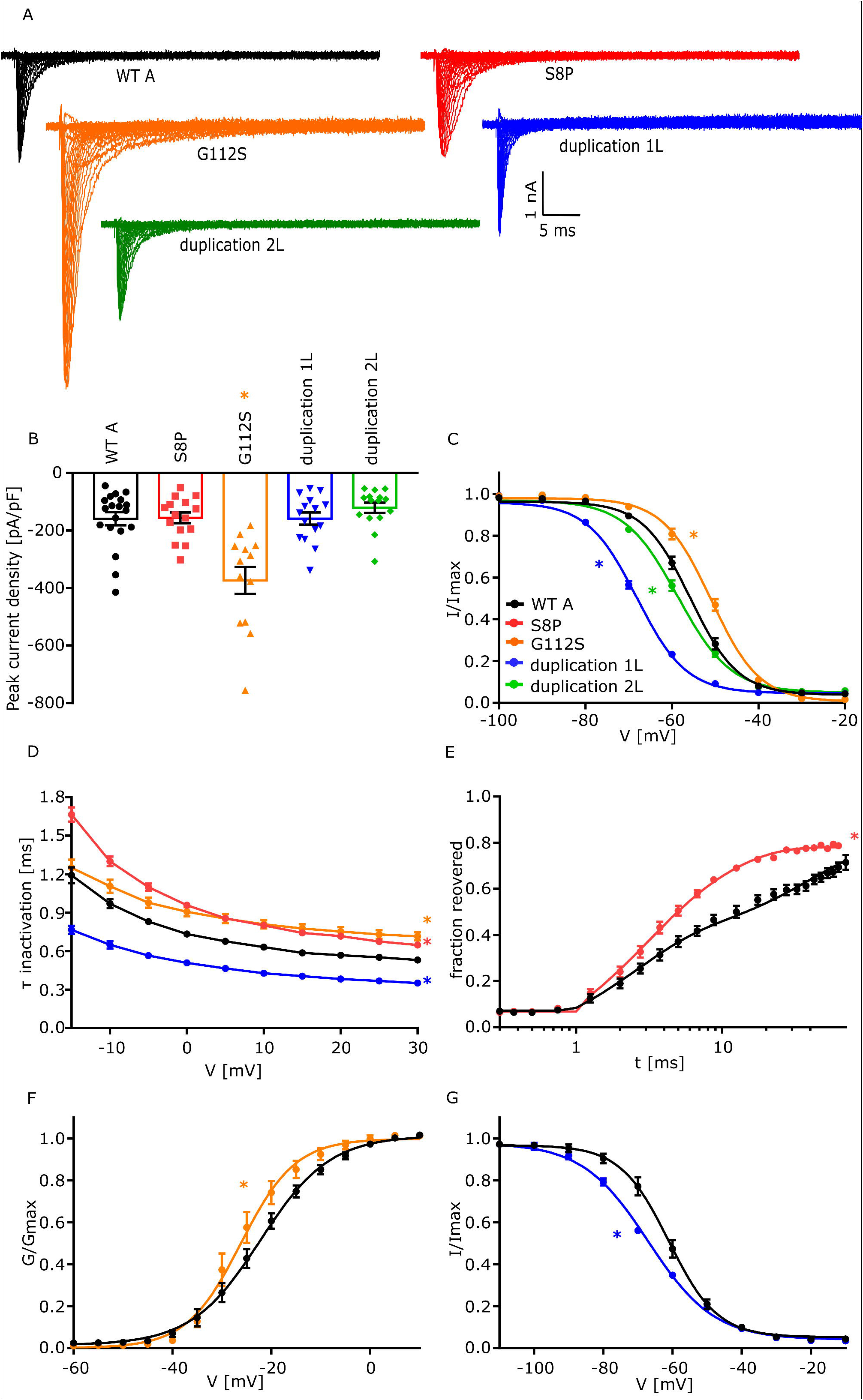
Electrophysiological analysis of FGF12 variants on Na_V_1.6. Functional analysis of the effect of 2 *FGF12* missense variants (S8P and G112S) and two *FGF12* CNVs (duplication 1L and duplication 2L) on Na_V_1.6 compared to WTA. In summary the figures show that the expression of both missense variants leads to a gain-of function in comparison to WTA while the two CNVs cause a loss-of function of the channel Na_V_1.6. A. Representative traces of Na_V_1.6 currents in ND7/23 cells expressing Na_V_1.6 with FGF12 WTA or the FGF12 variants, respectively, in response to voltage steps from -80 to +35 mV in 5 mV steps. B. Mean Current amplitudes of analysed ND7/23 cells injected with Na_V_1.6 with WTA (n=19), S8P (n=15), G112S (n=13), duplication 1L (n=15) or duplication 2L (n=15). Expression of Na_V_1.6 together with G112S showed a significantly elevated current density compared to Na_V_1.6 with WTA. C. Mean voltage-dependent fast inactivation of Na_V_1.6 with FGF12 WTA (n=16), G112S (n=10), duplication 1L (n=12) or duplication 2L (n=11). Lines illustrate Boltzmann Function fit to the data points. G112S showed a significant shift to more depolarized potentials in comparison to WTA while both Duplications show a hyperpolarizing shift in comparison to WTA. All data are shown as means ± SEM. D. Mean voltage-dependent fast inactivation time constant of Na_V_1.6 FGF12 WTA (n=19), S8P (n=15), G112S (n=13) or duplication 1L (n=15). S8P and G112S show a significantly slowed fast inactivation in comparison to WTA while duplication 1L shows an accelerated fast inactivation. All data are shown as means ± SEM. E. Time course of recovery from fast inactivation of Na_V_1.6 with FGF12 WTA (n=10) or FGF12 S8P (n=11) determined at -100 mV. The variant S8P leads to a significantly accelerated recovery of fast inactivation in comparison to WTA. Lines represent fits of 1 and τ2. A1 was set to 0.3. F. Mean voltage-dependent activation of Na_V_1.6 with FGF12 WTA (n=19) or G112S (n=13). G112S shows a hyperpolarizing shift in comparison to WTA. Lines illustrate Boltzmann Function fit to the data points. G. Mean voltage-dependent slow inactivation of Na_V_1.6 with FGF12 WTA (n=11) or FGF12 duplication 1L (n=10). Lines illustrate Boltzmann Function fit to the data points. FGF12 duplication 1L shows a shift to more hyperpolarized potentials in comparison to Na_V_1.6 with FGF12 WTA. All values of electrophysiological results, numbers and p-values are listed in table 1 and are shown as means ± SEM.

**Figure 4:**
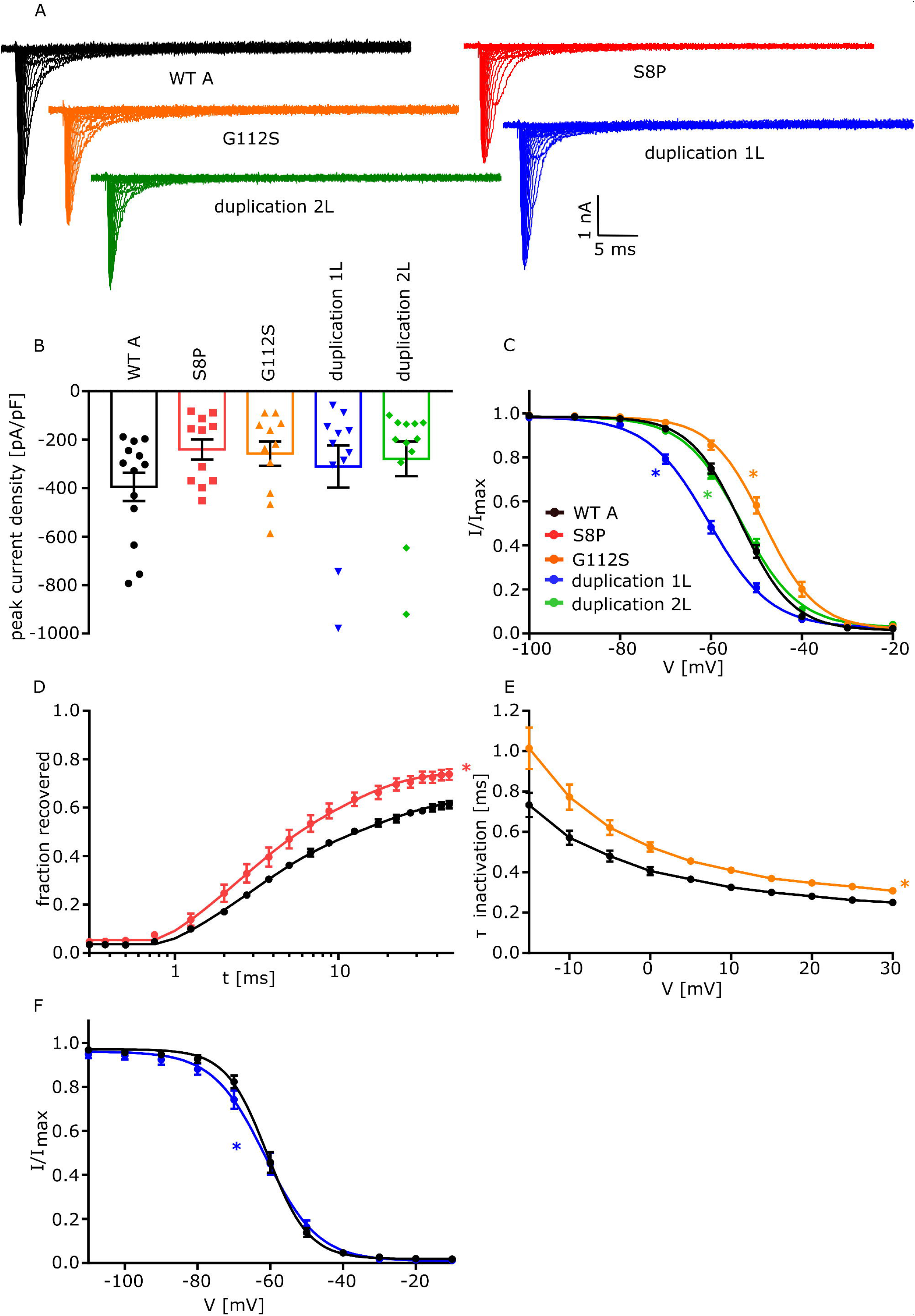
Electrophysiological analysis of FGF12 variants on Na_V_1.2. Functional analysis of the effect of 2 *FGF12* missense variants (S8P and G112S) and two *FGF12* CNVs (duplication 1L and duplication 2L) on Na_V_1.2 compared to WTA. In summary the figures show that the expression of both missense variants leads to a gain-of function in comparison to WTA while the two CNVs cause a loss-of function of the channel Na_V_1.2. A. Representative traces of Na_V_1.2 currents in ND7/23 cells expressing Na_V_1.2 with FGF12 WTA or the FGF12 variants, respectively, in response to voltage steps from -80 to +35 mV in 5 mV steps. B. Mean Current amplitudes of analysed ND7/23 cells injected with Na_V_1.2 with WTA (n=13), S8P (n=11), G112S (n=11), duplication 1L (n=11) or duplication 2L (n=12). None of the variants lead to a change in the current density of Na_V_1.2. C. Mean voltage-dependent fast inactivation of Na_V_1.2 with FGF12 WTA (n=13), G112S (n=11), duplication 1L (n=11) or duplication 2L (n=12). Lines illustrate Boltzmann Function fit to the data points. G112S showed a significant shift to more depolarized potentials in comparison to WTA while duplication 1L shows a hyperpolarizing shift in comparison to WTA and duplication 2L changes the slope of the fast inactivation. All data are shown as means ± SEM. D. Time course of recovery from fast inactivation of Na_V_1.2 with FGF12 WTA (n=11) or FGF12 S8P (n=10) determined at -100 mV. The variant S8P leads to a significantly accelerated recovery of fast inactivation in comparison to WTA. Lines represent fits of biexponential functions yielding the time constants τ1 and τ2. A1 was set to 0.3. E. Mean voltage-dependent fast inactivation time constant of Na_V_1.2 FGF12 WTA (n=11) or G112S (n=11). G112S show a significantly slowed fast inactivation in comparison to WTA. All data are shown as means ± SEM. F. Mean voltage-dependent slow inactivation of Na_V_1.2 with FGF12 WTA (n=10) or FGF12 duplication 1L (n=10). Lines illustrate Boltzmann Function fit to the data points. FGF12 duplication 1L shows a change in the slope of the slow inactivation in comparison to Na_V_1.2 with FGF12 WTA. All values of electrophysiological results, numbers and p-values are listed in table 2 and are shown as means ± SEM.

The second missense variant p.(Gly50Ser) (G50S) corresponds to WTB and is the same variant as p.(Gly112Ser) (G112S) which corresponds to WTA. This variant is located directly in the interaction interface between FGF12 and Na_V_1.6/Na_V_1.2 (Fig 1A). Co-expression of G112S with Na_V_1.6 led to an increased current density compared to WTA (Fig 3A and B, Table 1). Additionally, voltage-dependence of fast inactivation was shifted to more depolarized potentials, and fast inactivation was slowed (Fig 3C and D, Table 1). Furthermore, G112S showed a hyperpolarizing shift in the channel’s voltage dependence of activation, indicating a GOF effect when co-expressed with Na_V_1.6 (Fig 3F, Table 1). Co-expression of G50S together with Na_V_1.6 however only showed changes in fast inactivation, with a depolarizing shift of the voltage-dependence and a slowing of fast inactivation (Fig 5 A and B, Table 1) compared to WTB. A similar trend could be observed for G112S when expressed with the Na_V_1.2 channel. Again, a depolarizing shift of the voltage-dependence of fast inactivation and an additional slowing of fast inactivation led to a GOF of Na_V_1.2, comparable to a co-expression with the Na_V_1.6 channel (Fig 4C and E, Table 2). We noted a minor effect for the co-expression of G50S together with Na_V_1.2, with a change in just the slope factor of fast inactivation (Fig 5 D, Table 2).

**Figure 5:**
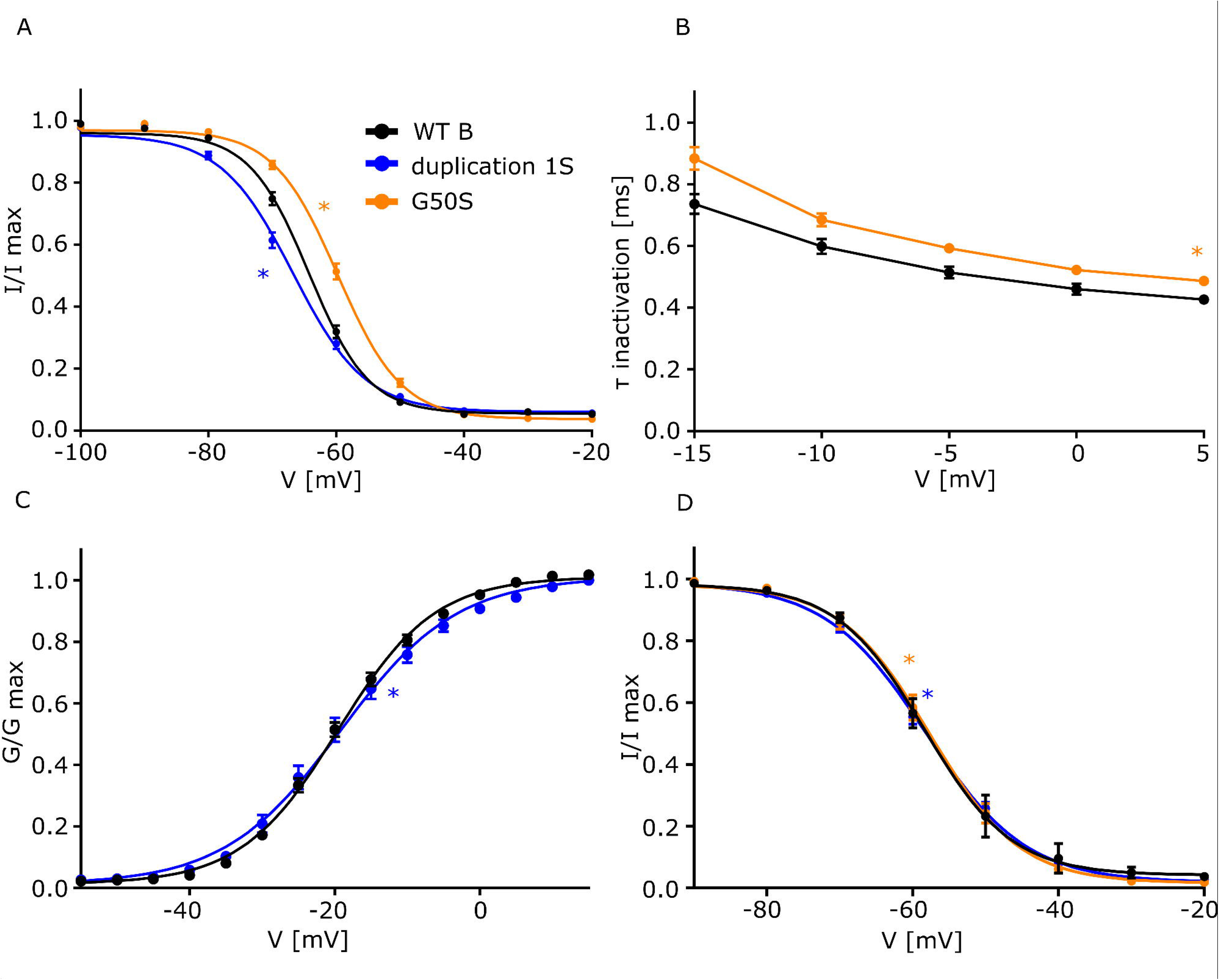
Electrophysiological analysis of FGF12 variants (NM_004113) on Na_V_1.6 or Na_V_1.2. Functional analysis of the effect of one *FGF12* missense variant (G50S) and one *FGF12* CNV (duplication 1S) on Na_V_1.6 and Na_V_1.2 compared to WTB. In summary the figures show that the expression of the missense variant leads to a gain-of function of Na_V_1.6 in comparison to WTB while the CNV cause a loss-of function of Na_V_1.6 in comparison to WTB. Duplication 2S did not show any significant effect on the channels. A. Mean voltage-dependent fast inactivation of Na_V_1.6 with FGF12 WTB (n=22), G50S (n=25) or duplication 1S (n=19). Lines illustrate Boltzmann Function fit to the data points. G50S showed a significant shift to more depolarized potentials in comparison to WTB while duplication 1S showed a hyperpolarizing shift in comparison to WTB. All data are shown as means ± SEM. B. Mean voltage-dependent fast inactivation time constant of Na_V_1.6 FGF12 WTB (n=22) or G50S (n=25). G50S showed a significantly slowed fast inactivation in comparison to WTB. All data are shown as means ± SEM. C. Mean voltage-dependent activation of Na_V_1.6 with FGF12 WTB (n=22) or duplication 1S (n=19). Duplication 1S shows a change in the slope of the activation curve in comparison to WTB. Lines illustrate Boltzmann Function fit to the data points. D. Mean voltage-dependent fast inactivation of Na_V_1.2 with FGF12 WTB (n=11), G50S (n=12) or duplication 1S (n=11). Lines illustrate Boltzmann Function fit to the data points. Both variants show a significant change in the slope of the fast inactivation in comparison to WTB. All data are shown as means ± SEM.

Interestingly, both duplication cases showed a completely different effect on Na_V_1.6 and Na_V_1.2 channels, compared to the effects of the missense variants detailed above. Since we used both transcript forms to analyse the different missense variants, we also used both forms to analyse the duplications. WTA without the last exon followed by exon 2, 3, 4 and 5 is subsequently referred to as duplication 1L, while WTB without the last exon followed by Exon 3, 4, 5 and 6 will be referred to as duplication 1S (Fig 1A and B). For duplication 2 we used WTA followed by WTB as duplication 2L, and WTB followed by WTB as duplication 2S (Fig 1A and C).

When co-expressing duplication 1L with Na_V_1.6, the voltage-dependence of fast inactivation was shifted to more hyperpolarized potentials, the time constant (τ) of fast inactivation was decreased and the voltage-dependence of slow inactivation was shifted to more hyperpolarized potentials when compared to WTA conditions (Fig 3C, D and G, Table 1), indicating a clear LOF. Co-expression of duplication 1S with Na_V_1.6 showed a hyperpolarizing shift of the fast inactivation curve and a change in the slope factor of the activation, leading to a LOF (Fig 5 A, C, Table 1). The co-expression of duplication1L with Na_V_1.2 showed similar effects as with Na_V_1.6, only without a change of the τ of fast inactivation (Fig 4C and F, Table 2). A similar trend was observed for the co-expression of duplication 1S with Na_V_1.2, which leads to a change in the slope factor in the fast inactivation curve (Fig 5 D). In summary, the expression of duplication 1 consistently leads to a LOF in both Na_V_1.6 and Na_V_1.2 channels.

The last variant, duplication 2L, only showed a hyperpolarizing shift of the voltage-dependence of fast inactivation when co-expressed with the Na_V_1.6 channel (Fig 3C, Table 1), while the co-expression of duplication 2S with Na_V_1.6 channel did not result in any effect on ion channel function. A similar trend could be observed for co-expression with Na_V_1.2. Here, duplication 2L only caused a minor change in the slope of the fast inactivation curve, while duplication 2S had no effect (Fig 4C, Table 2). Thus, duplication 2 gives rise to a LOF effect on both Na_V_1.6 and Na_V_1.2.

## Discussion

FGF12 had previously been suspected to be an important regulator of neuronal activity through modulation of different sodium channel subtypes.^5^ Here, we carried out a detailed analysis of the interaction of FGF12 with Nav1.2 and Nav1.6, which are known to be epilepsy-associated channels, and compared the physiological state with changes in ion channel function induced by missense and copy-number variants in *FGF12*.

Our functional analysis of WT FGF12 confirmed the gain-of-function (GOF) effect on Na_V_1.6 and the loss-of-function (LOF) effect on the voltage-dependent fast inactivation of Na_V_1.2, both of which have been described previously.^9, 12^ Furthermore, we characterized the differential effects of WT FGF12 on both channels in detail, providing evidence that FGF12 not just affects the voltage-dependence, but also the time constants of the fast inactivation of both channels. For Na_V_1.6, an additional effect on the voltage dependence of slow inactivation was evident.

Physiologically, FGF12 is expressed in two different transcripts resulting from alternative splicing, which differ only in their N-terminal part. Our results demonstrated that the expression of WTA, with its longer N-terminal part, has a stronger effect on the fast inactivation of Na_V_1.6 than WTB with its shorter N-terminal part, which is in line with previous studies.^9^ Additionally, the long N-terminal part of FGF12 WTA seems to affect the time constants of the fast inactivation, since WTB did not alter the time constants of fast inactivation. Thus, WTB caused a clear GOF of Na_V_1.6 channels, while WTA resulted in mixed GOF and LOF changes. The effect of the FGF12 WTs on Na_V_1.2 was less pronounced than for Na_V_1.6, but still reduced channel function.

Previously, missense variants or copy number variants (CNVs) in *FG12* have been solely identified in patients with DEE.^16^ Here we describe two additional patients, one with the recurrent variant G50S/G112S and a similar epileptic phenotype as described earlier^16^, and one patient with the novel variant S8P showing a completely novel phenotype consisting of autism spectrum disorder and developmental delay without seizures.

For sodium channels, like Na_V_1.6 and Na_V_1.2, it has already been shown that disease-causing variants can be associated with intellectual disability with or without seizures, depending primarily on the resulting kinetic changes within the channel.^3, 24^ LOF variants in Na_V_1.6 have mainly been associated with intellectual disability, while GOF variants lead to an epileptic phenotype. For Na_V_1.2, a similar genotype-phenotype correlation has been observed, with the difference that Na_V_1.2 LOF caused by missense variants can also cause severe late onset epilepsy phenotypes, while truncating variants (which by definition cause complete LOF) are associated with ASD.^25^

Interestingly, the functional analysis of variant S8P, which was found in a patient with ASD, caused GOF changes in Na_V_1.6 and Na_V_1.2 channels, mainly by affecting the time constant of fast inactivation components of both channels. These changes lead to a stronger deceleration of fast inactivation of Na_V_1.6 and reduced the regulating effect of WTA on the recovery from fast inactivation. Additionally, we could show that the effect on the recovery from fast inactivation of FGF12 WTA on Na_V_1.2 was reduced as well, which results in a GOF for both channels. This is an unexpected finding, since ASD has so far primarily been associated with LOF of Na_V_1.6 or Na_V_1.2 channels.^3, 25^

For the second missense variant G50S/G112S (WTB/WTA variants), only two patients have been described before. Here, we identified an additional patient with this variant showing a similar age of onset at about three months of age, and phenotypic features including generalized and focal seizures and moderate intellectual disability. This variant seems to influence Na_V_1.6 in both forms in a similar way leading to a 5 mV depolarizing shift in the voltage-dependence of fast inactivation as well as to a decelerated fast inactivation. The same effect is visible for G112S on Na_V_1.2 while G50S shows just a small effect on the voltage-dependence of fast inactivation on Na_V_1.2 but does not influence the time constant of fast inactivation. In summary, the gain-of function effect of the variant G50S/G112S on Na_V_1.6 on the voltage-dependence of fast inactivation was similar to the effect of the already described variant R52H/R114H on Na_V_1.6^9^, which is located just two amino acids apart in the interaction interface of FGF12 to the Na_V_s. The similar effect on the voltage-dependence of fast inactivation and the very close localization of these two variants suggests that both mutated FGF12 proteins may potentially interfere with channel function through a similar mechanism.

Analysis of the effect of CNVs on Na_V_1.6 and Na_V_1.2 revealed a key difference when compared to the effect of the missense variants R52H/R114H and G50S/G112S. Both CNVs primarily affected the channels’ fast inactivation, leading to a clear LOF of both channels. One possible explanation may be that the duplication 1L partially lost the ability to interact with Na_V_1.6, since the inactivation kinetics were similar to that of Na_V_1.6 expressed without FGF12. Only the effect on the peak current density and the recovery from fast inactivation were comparable to co-expressed WTA and Na_V_1.6. In contrast, duplication 1L seems to negatively regulate Na_V_1.2 since fast inactivation, which is not changed by WTA, showed a hyperpolarizing shift during co-expression with duplication 1L. This interaction also appeared to affect the slow inactivation, which was unchanged by wildtype co-expression. Further experiments are needed to analyse if the interaction between the CNVs and the channels is weakened or changed in another way, since some of the kinetic parameters which are changed by WTA are not changed anymore by the CNVs.

Interestingly, both duplications caused a similar phenotype regarding seizure types, age of onset and moderate to severe developmental delay/intellectual disability. This overlaps with the symptomatic spectrum associated with the missense variants R52H/R114H and G50S/G112S. The only difference between the point mutations and the CNVs was the age of onset, with later onset in the duplication cases at a median age of onset of 15.5 months (n = 4).^19, 23^ This finding is in line with previous studies, where Na_V_1.2 LOF variants were shown to be associated with a later disease onset.^25^

In summary, the missense variant G50S/G112S led to a clear GOF effect on both channels, while both duplications were associated with LOF effects. The phenotype of the patients with these variants showed significant overlap to the established genotype-phenotype spectrum of GOF and LOF variants in *SCN2A*, but patients with LOF variants are known to show severe DEE with a later age of onset.^25^ Thus, no clear genotype-phenotype correlation was visible for these variants. Surprisingly, the variant S8P led to a GOF effect on both channels, which is a functional effect that has not been previously described for ASD associated with variants in sodium channel genes.

In a previous study, 6 out of 17 patients carrying one of the already published *FGF12* variants showed a partial response to phenytoin or lamotrigine, sodium channel blockers that enhance the channels’ fast inactivation by shifting voltage-dependence towards more hyperpolarized potentials.^16^ However, not all patients responded well to phenytoin, suggesting that these variants may have an additional effect which could not be identified until now. This hypothesis is supported by the FGF12 mouse variant model *fgf12*^R52H/+^ which suffers from spontaneous seizures and dies of sudden unexplained death in epilepsy (SUDEP) at around 16 days of age.^26^ Veliskova and co-workers treated these mice with phenytoin, but the survival time was just marginally prolonged to about 19 days.^26^

In this study, we were the first to investigate the detailed biophysical and physiological modification of the channel function of Nav1.2 and Nav1.6 by wildtype FGF12. We found a divergent effect on the sodium channel function, with a trend towards GOF in Nav1.6 and a trend towards LOF in Nav1.2. Secondly, we described the underlying pathomechanism of four *FGF12* missense and copy-number variants leading to complex gain- or loss-of function changes in Na_V_1.6 and Na_V_1.2. Lastly, we identified two new patients - one with DEE and one with ASD - expanding the phenotypic spectrum of FGF12-associated disorders to include ASD, which has not been previously described. This establishes FGF12 as an important modulator of neuronal activity driven by sodium channel subtypes and confirms its role as an epilepsy gene with an overlap to intellectual disability and autism spectrum disorder.

## Acknowledgement

Y.W. was supported by the DFG/FNR INTER Research Unit FOR2715 of the German Research Foundation (We4896/4-1 and WE4896/4-2) and by the BMBF Treat-ION grant (01GM1907).

I.H. received support through the German Research Foundation (HE5415/6-1) and the DFG/FNR INTER Research Unit FOR2715 (He5415/7-1 and He5415/7-2). I.H. was supported by The Hartwell Foundation through an Individual Biomedical Research Award. This work was also supported by the National Institute for Neurological Disorders and Stroke (K02 NS112600), including support through the Center Without Walls on Ion channel function in epilepsy (“Channelopathy-associated Research Center”, U54 NS108874), the Eunice Kennedy Shriver National Institute of Child Health and Human Development through the Intellectual and Developmental Disabilities Research Center (IDDRC) at Children’s Hospital of Philadelphia and the University of Pennsylvania (U54 HD086984), and by intramural funds of the Children’s Hospital of Philadelphia through the Epilepsy NeuroGenetics Initiative (ENGIN). Research reported in this publication was also supported by the National Center for Advancing Translational Sciences of the National Institutes of Health under Award Number UL1TR001878. The content is solely the responsibility of the authors and does not necessarily represent the official views of the NIH. This project was also supported in part by the Institute for Translational Medicine and Therapeutics’ (ITMAT) Transdisciplinary Program in Translational Medicine and Therapeutics at the Perelman School of Medicine of the University of Pennsylvania. I.H. also received support through the International League Against Epilepsy (ILAE). The study also received support through the EuroEPINOMICS-Rare Epilepsy Syndrome (RES) Consortium and by the Genomics Research and Innovation Network (GRIN, grinnetwork.org).

## Author contribution

SS performed genetic and functional analysis and wrote the manuscript together with YW

MP, TB and AvdV performed detailed clinical evaluation and further genetic analysis and edited the manuscript

NS helped to conceptualize the used constructs for the electrophysiological measurements and edited the manuscript

HG prepared cDNA samples from venous blood from the patients and edited the manuscript

MW prepared fibroblasts from a skin biopsy of the patient and edited the manuscript

UBSH re-analyzed the functional data and edited the manuscript

IH planned the study together with YW, supervised the analysis and edited the manuscript.

CMB reviewed the results and edited the manuscript.

YW planned and coordinated the study, supervised the analysis, wrote, and edited the manuscript together with SS

## References

1. Catterall, W. A. Voltage-gated sodium channels at 60: structure, function and pathophysiology: Voltage-gated sodium channels. J. Physiol. 590, 2577–2589 (2012).

2. Brunklaus, A., Ellis, R., Reavey, E., Semsarian, C. & Zuberi, S. M. Genotype phenotype associations across the voltage-gated sodium channel family. J. Med. Genet. 51, 650–658 (2014).

3. Liu, Y. et al. Neuronal mechanisms of mutations in *SCN8A* causing epilepsy or intellectual disability. Brain 142, 376–390 (2019).

4. Zhang, S. et al. SCN9A Epileptic Encephalopathy Mutations Display a Gain-of-function Phenotype and Distinct Sensitivity to Oxcarbazepine. Neurosci. Bull. 36, 11–24 (2020).

5. Goldfarb, M. et al. Fibroblast Growth Factor Homologous Factors Control Neuronal Excitability through Modulation of Voltage-Gated Sodium Channels. Neuron 55, 449–463 (2007).

6. Smallwood, P. M. et al. Fibroblast growth factor (FGF) homologous factors: new members of the FGF family implicated in nervous system development. Proc. Natl. Acad. Sci. 93, 9850–9857 (1996).

7. Laezza, F. et al. FGF14 N-terminal splice variants differentially modulate Nav1.2 and Nav1.6-encoded sodium channels. Mol. Cell. Neurosci. 42, 90–101 (2009).

8. Yang, J. et al. FGF13 modulates the gating properties of the cardiac sodium channel Na _v_ 1.5 in an isoform-specific manner. Channels 10, 410–420 (2016).

9. Siekierska, A. et al. Gain-of-function *FHF1* mutation causes early-onset epileptic encephalopathy with cerebellar atrophy. Neurology 86, 2162–2170 (2016).

10. Wildburger, N. C. et al. Quantitative Proteomics Reveals Protein–Protein Interactions with Fibroblast Growth Factor 12 as a Component of the Voltage-Gated Sodium Channel 1.2 (Nav1.2) Macromolecular Complex in Mammalian Brain*. Mol. Cell. Proteomics 14, 1288–1300 (2015).

11. Liu, C., Dib-Hajj, S. D., Renganathan, M., Cummins, T. R. & Waxman, S. G. Modulation of the Cardiac Sodium Channel Nav1.5 by Fibroblast Growth Factor Homologous Factor 1B. J. Biol. Chem. 278, 1029–1036 (2003).

12. Wang, C., Wang, C., Hoch, E. G. & Pitt, G. S. Identification of Novel Interaction Sites that Determine Specificity between Fibroblast Growth Factor Homologous Factors and Voltage-gated Sodium Channels. J. Biol. Chem. 286, 24253–24263 (2011).

13. Dalski, A. et al. Mutation analysis in the fibroblast growth factor 14 gene: frameshift mutation and polymorphisms in patients with inherited ataxias. Eur. J. Hum. Genet. 13, 118–120 (2005).

14. Puranam, R. S. et al. Disruption of Fgf13 Causes Synaptic Excitatory-Inhibitory Imbalance and Genetic Epilepsy and Febrile Seizures Plus. J. Neurosci. 35, 8866–8881 (2015).

15. Al-Mehmadi, S., et al. FHF1 (FGF12) epileptic encephalopathy: Table. Neurol. Genet. 2, e115 (2016).

16. Trivisano, M. et al. Defining the phenotype of *FHF1* developmental and epileptic encephalopathy. Epilepsia 61, (2020).

17. Sobreira, N., Schiettecatte, F., Valle, D. & Hamosh, A. GeneMatcher: A Matching Tool for Connecting Investigators with an Interest in the Same Gene. Hum. Mutat. 36, 928–930 (2015).

18. Richards, Sue et al. Standards and guidelines for the interpretation of sequence variants: a joint consensus recommendation of the American College of Medical Genetics and Genomics and the Association for Molecular Pathology. Genet. Med. 17, 405–423 (2015).

19. Willemsen, M. H. et al. Epilepsy phenotype in individuals with chromosomal duplication encompassing *FGF12*. Epilepsia Open 5, 301–306 (2020).

20. Leffler, A., Herzog, R. I., Dib-Hajj, S. D., Waxman, S. G. & Cummins, T. R. Pharmacological properties of neuronal TTX-resistant sodium channels and the role of a critical serine pore residue. Pflüg. Arch. - Eur. J. Physiol. 451, 454–463 (2005).

21. Rush, A. M., Dib-Hajj, S. D. & Waxman, S. G. Electrophysiological properties of two axonal sodium channels, Na _v_ 1.2 and Na _v_ 1.6, expressed in mouse spinal sensory neurones: Sodium channels in sensory neurones. J. Physiol. 564, 803–815 (2005).

22. Wood, J. N. et al. Novel cell lines display properties of nociceptive sensory neurons. Proc. R. Soc. Lond. B Biol. Sci. 241, 187–194 (1990).

23. Shi, R.-M. et al. Phenytoin-responsive epileptic encephalopathy with a tandem duplication involving FGF12. Neurol. Genet. 3, e133 (2017).

24. Ben-Shalom, R. et al. Opposing Effects on Na V 1.2 Function Underlie Differences Between SCN2A Variants Observed in Individuals With Autism Spectrum Disorder or Infantile Seizures. Biol. Psychiatry 82, 224–232 (2017).

25. Wolff, M. et al. Genetic and phenotypic heterogeneity suggest therapeutic implications in SCN2A-related disorders. Brain 140, 1316–1336 (2017).

26. Velíšková, J. et al. Early onset epilepsy and sudden unexpected death in epilepsy with cardiac arrhythmia in mice carrying the early infantile epileptic encephalopathy 47 gain of function *FHF1(FGF12)* missense mutation. Epilepsia 62, 1546–1558 (2021).

